# Dynamic dorsal body morphology encodes engineering design principles of fish propulsion and hydrodynamics

**DOI:** 10.64898/2026.05.06.723159

**Authors:** Yunpeng Zhu, Lidong Zhu, Lianyuan Cheng, Liangliang Cheng, Xingwen Zheng, Duncan Irschick, Johnson Martin, J. Nathan Kutz

**Affiliations:** School of Engineering and Materials Science, Queen Mary University of London, London, UK; Institute of Cyber-Systems and Control, College of Control Science and Engineering, Zhejiang University, Hangzhou 310027, China; ENTEG, Faculty of Science and Engineering, University of Groningen, Groningen, 9747 AG, The Netherlands; Institute of Fundamental and Transdisciplinary Research, Zhejiang University, Hangzhou, 310058, China; State Key Laboratory of Ocean Sensing, Zhejiang University, Hangzhou, 310058, China; Department of Biology, University of Massachusetts at Amherst, Massachusetts, USA; 2329 E Main St Unit 3, Wilmore, Kentucky, USA; Autodesk Research, 6 Agar Street, London, UK; Department of Applied Mathematics and Electrical and Computer Engineering, University of Washington, Seattle, WA USA

**Keywords:** Inverse design, biological locomotion, fluid-structure interaction, machine learning

## Abstract

Understanding how biological shape and movement interact with surrounding fluids represents a fundamental challenge at the intersection of biology, physics, and engineering. Fish locomotion exemplifies this challenge: body morphology and swimming kinematics together determine the hydrodynamic forces and flow structures that enable efficient propulsion and maneuverability. Whereas biologists have long sought to connect morphological variation to swimming performance, traditional morphometric approaches provide limited insight into the fluid mechanical consequences of shape differences. Similarly, although computational fluid dynamics can reveal detailed flow physics, simulating hydrodynamics across diverse and dynamic morphologies remains prohibitively expensive for systematic investigation. To bridge this gap, we introduce a data-driven framework that connects fish body shape dynamics to hydrodynamic performance through compact morphospace parameterization and reduced-order modeling. Using CFD simulations of 15 fish species from the Digital Life Project database (www.digitallife3d.org/3d-model), we generate hydrodynamic datasets capturing the shape-flow relationship. Principal Component Analysis (PCA) extracts four dominant shape parameters from dorsal body profiles, which are then integrated into an Inverse-Design with Dynamic Mode Decomposition (ID-DMD) framework to model the resulting fluid dynamics. The resulting modal analysis suggests that locomotion strategies emerge from specific shape-flow interactions. We further demonstrate the framework’s utility through single- and multi-objective shape optimization, showing how it enables efficient exploration of the morphology-hydrodynamics relationship. This approach offers a novel analysis and design tool for understanding how biological form and motion interact with fluid mechanics, with applications ranging from bio-inspired vehicle development to evolutionary biomechanics.

---

The extraordinary diversity of fish body shapes reflects millions of years of adaptation to life in moving fluids. Despite their evolutionary divergence, all fish face the same fundamental physical constraints of swimming: generating thrust and maneuverability while minimizing drag and maintaining stability [1, 2, 3, 4, 5]. Identifying the design principles that link body morphology to locomotor function has therefore long been a central goal in biomechanics, evolutionary biology, and bio-inspired engineering [6, 7, 8, 9, 10]. Existing morphological analyzes typically rely on previously hypothesized form/function relationships, which may lack the required level of detail to deeply understand how body shape may impact hydrodynamic performance. This is an especially important issue given the very large diversity of fish species (*>* 30, 000), and body shapes [11, 12] or purely statistical descriptors, which fail to capture subtle, spatially distributed shape variations that govern hydrodynamic performance [13, 14, 5]. On the other hand, studies using high-fidelity Computational Fluid Dynamics (CFD), while extremely valuable, are prohibitively expensive and time-consuming for systematic exploration across species and morphologies for hydrodynamic properties [15, 16, 17, 18]. Here, we introduce a data-driven framework that addresses these issues, which integrates diverse fish body shapes into a continuous latent morphospace. By representing morphology as a smooth, hydrodynamic-relevant manifold, this approach enables the understanding of evolutionary design rules while directly supporting the inverse design and optimization of bio-inspired swimming systems.

The functional mapping between morphology and locomotion is central to biological diversification, as body shape must balance competing demands for maneuverability, speed, and energetic economy [2, 19, 20, 21, 22]. Yet, existing morphometric tools struggle to fully capture this complexity. Traditional distance-based approaches are easy to interpret but reduce continuous biological shapes to simple morphometric ratios, often missing the subtle geometric features that are critical to hydrodynamic interaction [23, 24]. Landmark-based Geometric Morphometrics (GM) captures spatial structure, yet its reliance on homologous landmarks makes it difficult to describe the smooth outlines and deformable regions essential for swimming [14, 25, 26]. While outline-based methods like the Elliptic Fourier Transform (EFT) can recover global geometry [27, 28, 29], their coefficients remain mathematically abstract. This lack of direct morphological meaning makes it difficult to relate shape variation back to functional performance. Consequently, researchers are often forced to choose between geometric precision and biological interpretability, leaving the morphological drivers of locomotive evolution poorly understood.

While high-fidelity CFD can explicitly bridge morphology to hydrodynamic performance, its prohibitive computational cost due to accurately resolving the dynamic morphology and nuanced flow features precludes systematic exploration across broad aquatic morphospaces [17]. Recent advances in machine learning have introduced surrogate modeling as a scalable alternative, utilizing operator-learning frameworks, such as Deep Operator Networks (DeepONets) and Fourier Neural Operators (FNO), to approximate complex flow solutions [30, 31, 32]. These methods can efficiently map geometric parameters to flow fields, yet they often rely on highly parameterized model architectures that require extensive training data and time [33]. Furthermore, their performance on complex spatio-temporal scientific data does not meet the necessary standards of accuracy [34]. Their “black-box” nature also obscures the underlying physical principles, limiting their ability to yield the mechanistic insights essential for biological discovery or bio-inspired engineering.

Leveraging high-resolution body geometries from 15 divergent fish species in the Digital Life Project [35], we employ *Principal Component Analysis* (PCA) to extract the dominant modes of morphological variation (fish body shapes). The PCA analysis is performed using fish specimens that are posed in a neutral (non-bent) position. Specifically, the PCA reveals that four modes dominate the morphospace across the species considered, with the variance of PCA modes two, three, and four determining key elements between fish that possess morphological shapes for powerful thrust mechanics versus those species who exhibit morphological shapes for energy efficient cruising. These dominant PCA modes allow for the construction of a morphospace that accurately characterizes all fish shapes studied. Moreover, it allows for a low-dimensional representation that is ideal for inverse design and the construction of reduced-order models of fish propulsion.

Unlike traditional approaches, the PCA formulation embeds shape parameters directly within a dynamic modeling process, allowing morphology to be systematically explored and optimized within a unified, continuous manifold. We integrate these morphological modes into an Inverse-Design Dynamic Mode Decomposition (ID-DMD) framework [36], which preserves the physical interpretability of modal analysis (e.g., spatial structures, damping rates, and oscillation frequencies) while delivering predictions orders of magnitude faster than direct numerical simulation (Fig.1). The analysis reveals a direct correspondence between dorsal fish body geometry and wake vortex topologies, demonstrating that divergent swimming strategies are intrinsically encoded in these morphological modes. By synthesizing biological form with fluid mechanics, this framework translates the underlying dynamics of aquatic locomotion into engineering design principles, informing the development of next-generation bio-inspired autonomous systems.

**Figure 1:**
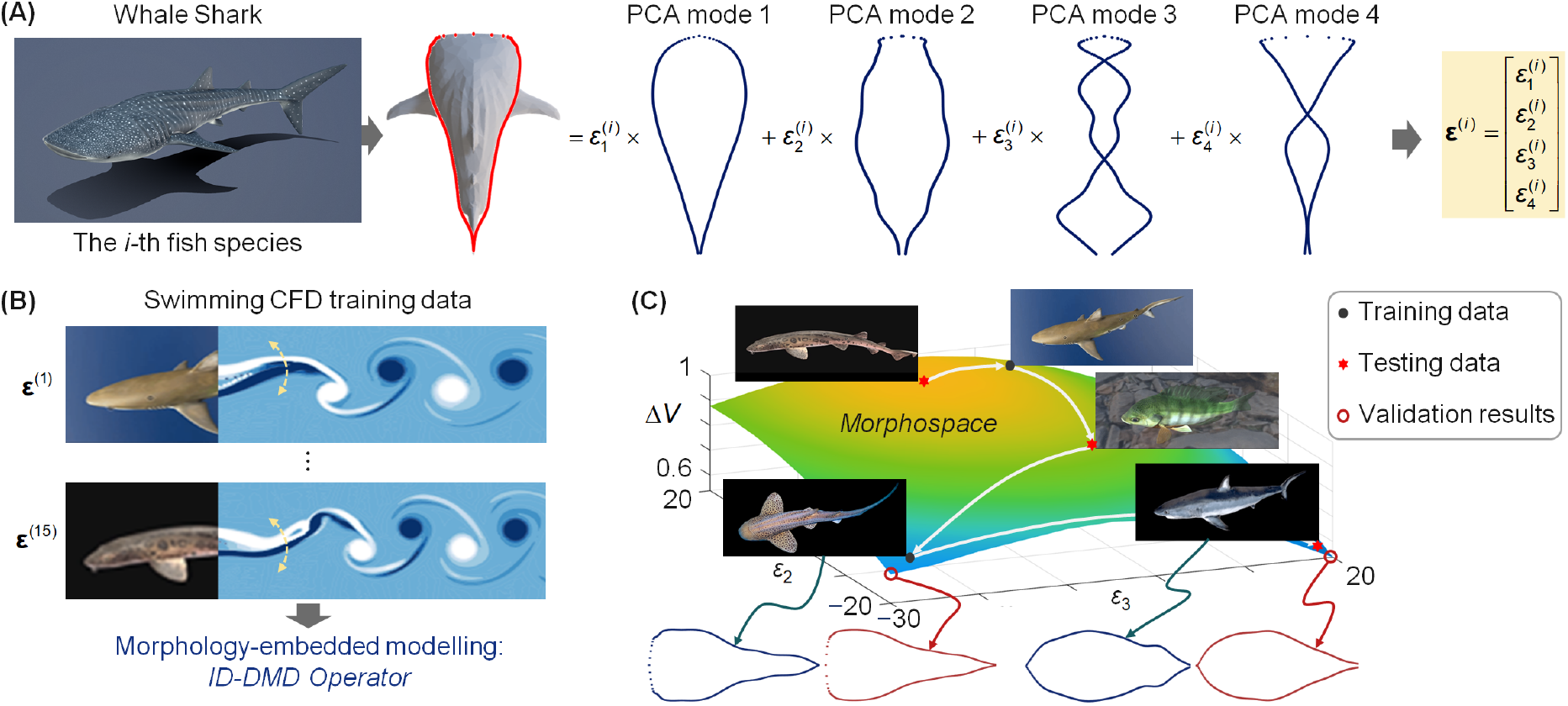
The data-driven framework for dorsal fish-body morphological parameterization and fluid dynamic design. **(A).** Biological fish morphology illustrated using a Whale Shark, together with its dorsal-view geometry, where the fish body shape are parameterized using a set of PCA modes which form a morphospace, where the coefficients *ε*_1_, *ε*_2_, *ε*_3_, *ε*_4_ control approximately 90% of the variations in body geometries. **(B)**. CFD simulations of various fish morphologies while swimming, controlled by **ε** = {*ε*_1_, *ε*_2_, *ε*_3_, *ε*_4_} produce comprehensive hydrodynamic datasets, such as Vortex shedding, Field pressure, and induced velocity. An ID-DMD model is directly identified from these CFD simulation data, embedding the morphological parameters **ε** into the dynamical model to predict local body motion and the associated flow fields. **(C)**. The construction of the morphospace is a smooth, low-dimensional design space (i.e., the dynamic range of vorticity of the local motion (Δ*V*) against the second- and third-order shape mode coefficients, with other coefficients fixed) for efficient analysis and optimization of bio-inspired fish-body shapes. Note that the morphospace correctly predicts held-out test data, which represent new fish and their body shapes.

## Results

### Parameterization of Fish Body Geometries

We applied high-resolution 3D scans of 15 living fish species developed in the Digital Life Project [35]. This dataset — which includes the apex-predatory Shortfin Mako to the sedentary Epaulette Shark—represents a broad spectrum of aquatic design solutions as listed in Table.1. Indeed, the data ranges from the Largemouth Bass which has an average length and weight of 18 inches and 3 pounds respectively, to the Whale Shark which is on average 40 feet and 11 tons. Thus there is more than an order of magnitude difference in length and five orders of magnitude difference in weight represented in the data set. These 3D models were created to accurately represent the generic body shapes of the actual fish species for a given age and sex, as described in detail in several research studies [37, 38, 39].

**Table 1:**
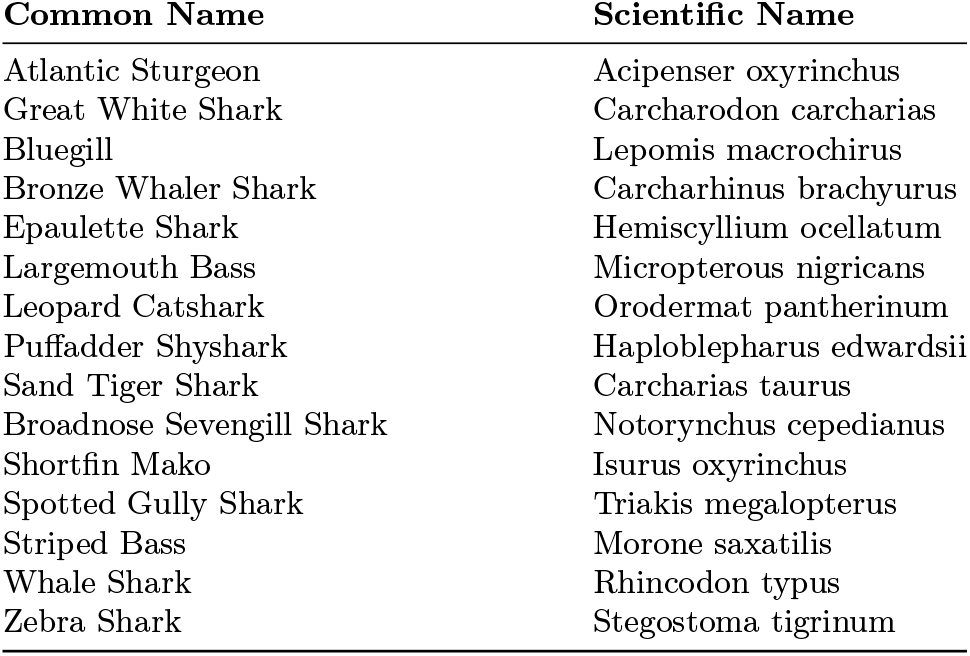
List of the 15 fish species used in this study.

In order to embed this morphological complexity into a continuous and tractable design space, we applied PCA to the dorsal-view body shapes. For the PCA, the body shapes were in a neutral straight pose which is normalized to be of the same length and temporal swimming frequency (See *Supplementary Information Section I and II*). The results indicate that the vast morphological diversity of the 15 fish species can be efficiently compressed into a linear combination of four dominant shape modes (Fig.1 (B)). The leading mode has a nearly constant coefficient (*ε*_1_) across species and captures the baseline body geometry of all, i.e. wide at the head and tapering towards the tail. The subsequent PCA coefficients (**ε**_d_ = {*ε*_2_, *ε*_3_, *ε*_4}_) capture the shape variations of fish body geometries (*Fig.S1*) that have distinct hydrodynamic performances designed by nature. This formulation maps discrete biological specimens onto a continuous low-dimensional latent morphospace spanned by these variable PCA coefficients, enabling systematic interpolation and generation of new fish geometries in the morphospace. The morphological parameterization process is introduced in Materials and Methods, see detailed justifications in *Supplementary Information Section I*.

### Fish Hydrodynamics on a Unified Morphospace

To bridge the gap between fish morphology and hydrodynamic function (i.e. swimming fish), we integrated the PCA-derived morphospace into an ID-DMD framework (Fig.1 (C)) [36]. ID-DMD is based on simple least-square regression and the underlying dynamic mode decomposition (DMD) algorithm [40] which can be used to construct a low-rank subspace spanning multiple experiments in parameter space. The ID-DMD algorithm leverages the computed low-dimensional subspace to enable fast digital design and optimization on laptop-level computing, including the potential to prescribe the dynamics themselves. Moreover, the method is robust to measurement noise, physically interpretable to key dynamic traits, such as system stability (damping) and rhythmic cycles (frequencies), and can provide uncertainty quantification metrics [41]. The ID-DMD architecture can also efficiently scale to large-scale design problems using randomized algorithms [42]. The simplicity of the method and its implementation are highly attractive in practice, and the ID-DMD has been demonstrated to be an order of magnitude more accurate than competing methods while simultaneously being 3-5 orders faster on challenging engineering design problems ranging from structural vibrations and fluid dynamics to real-world experimental droplet control [36].

Mathematically, we consider the state space of our system at time *t*_*k*_ to be given by **x**(*t*_*k*_) = **x**_*k*_. The state space in our case is a column vector of all the flow field properties computed in simulations of the swimming motion (e.g., vorticity, velocity, and pressure). Although the fields are two-dimensional from a dorsal plane view, they are flattened and stacked to produce the overall state-space at a given time point. This is then arranged as a temporal sequence model (input-output model)

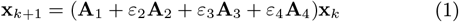

where **A**_*i*_, *i* = 1, 2, 3, 4 are learned operator matrices with constant entries that map the state space from time *t*_*k*_ to *t*_*k*+1_. ID-DMD explicitly assumes that the dynamics over a time step can be approximated by linear mappings represented by the **A**_*i*_, where the parameters *ε*_*i*_ explicitly encode the dependence on the PCA body shape modes. Standard DMD does not allow for dependencies on the parameters *ε*_*i*_. Further, we note that the dependence on the base body shape parametrized by *ε*_1_ is not included, given that this shape has essentially no variability between fish species. Even for exceptionally complex flow fields, the DMD approximation can be an accurate approximation to the underlying nonlinear dynamics. Indeed, DMD was introduced in the CFD community by Schmid [43] who presented it as a tool for the modal analysis of complex flow fields. DMD has since been used successfully across a large number of domains [40] and as an effective parametric model reduction paradigm [44, 45, 46].

To learn the ID-DMD model (7), the ID-DMD algorithm is applied to CFD simulations whose parameters and settings (*Fig.S2*) are introduced in *Supplementary Information Section II*. The ID-DMD model captures the dynamics of vortex shedding, the fluid velocity fields, and the corresponding pressure distributions. The accuracy of the model predictions against high-fidelity CFD simulations is validated in Fig.2, where the relative error is defined as

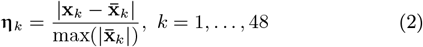

where 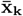 is the predicted ID-DMD model response (See *Table S2* for all testing results).

**Figure 2:**
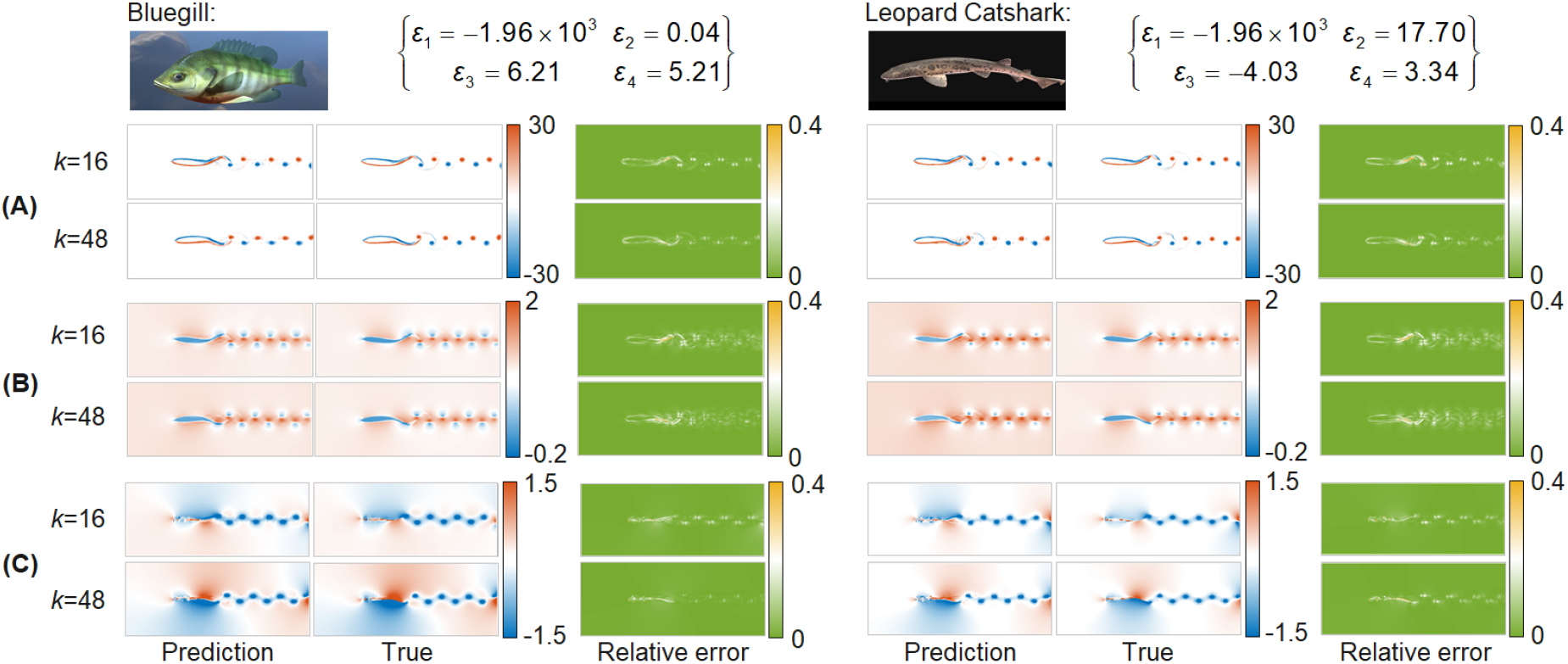
The Prediction of local hydrodynamics for fish species not included in the training data, i.e. Bluegill and Leopard Catshark, using the ID-DMD model and relative errors at different time steps (*k*=16 and 48), where the maximum error of approximately 0.4 occurs at the fluid-structure boundary. **(A).** Prediction of vortex shedding patterns: A vortex is defined as a region of rotating fluid flow, where the structure and strength of these shed vortices represent how much thrust a fish is producing and how efficiently it is swimming. **(B)**. Prediction of induced velocity fields: The induced velocity field is created when the tail pushes against the water, which directly links to propulsive efficiency. An efficient swimmer is a master at keeping induced velocity low relative to the thrust it produces. **(C)**. Prediction of pressure distributions: Pressure is the mechanism of force generation, where a low-pressure suction zone facilitates forward motion by pulling the fish through the fluid. The accurate prediction of these hydrodynamic patterns demonstrates that the embedded morphological parameters **ε** effectively govern fluid-structure interactions across a diverse range of geometries within the morphospace.

The ID-DMD approach demonstrates orders of magnitude faster prediction than CFD simulations and existing advanced machine learning approaches, as discussed in the Materials and Methods. More importantly, the modal interpretability of ID-DMD enables hydrodynamic responses to be directly linked to specific morphological changes (*Fig.S3*) as discussed in Materials and Methods and *Supplementary Information Section III*.

### Optimized Body Shape for Fish Propulsion

The latent morphospace generated by a PCA decomposition serves as a lowdimensional manifold for fish shape optimization. Here, we conducted single- and multi-objective optimizations of propulsion properties based on the ID-DMD model, where the optimization problems are introduced in Materials and Methods. This process effectively allows the model to navigate the manifold of possible fish shapes to locate the global optima governed by fluid mechanical constraints.

The single-objective optimization aims to minimize the normalized range of vorticity of the hydrodynamic flow of fish bodies, a proxy for the propulsive intensity of the wake structures as illustrated in Fig.1 (C). In the context of high-speed cruising, a reduced vorticity range signifies a minimized shear layer loss and a more streamlined energy transfer from the body to the wake. It is clear that the body shapes reconstructed from the two minimum points are very close to the actual 3D model, The Shortfin Mako shark and Zebra Shark, respectively, have the lowest vorticity shifts among all the 15 fish species as illustrated in *Fig.S4*. In addition, multi-objective optimization is conducted to demonstrate the versatility of the ID-DMD approach in handling multiple flow quantities that are critical to hydrodynamic performance. The design results are shown in Fig.3 by minimizing the dynamic range of vorticity, induced velocity, and flow field pressure around a fish body [47, 48], where the high-propulsive efficiency (the Froude efficiency) design targets converge toward the morphological characteristics of the Shortfin Mako, which is a very fast species with powerful thrust performance [49]. Non-optimized baselines, such as the Zebra Shark, are characterized by lower propulsive efficiency but higher locomotor flexibility. This is consistent with the ecology of Zebra sharks, which have evolved for slower-speed cruising [50]. This bifurcation in the Pareto front suggests that our model accurately captures the fundamental trade-offs between energetic economy, maneuverability, and speed inherent in aquatic locomotion. Details are discussed in *Supplementary Information Section IV*.

**Figure 3:**
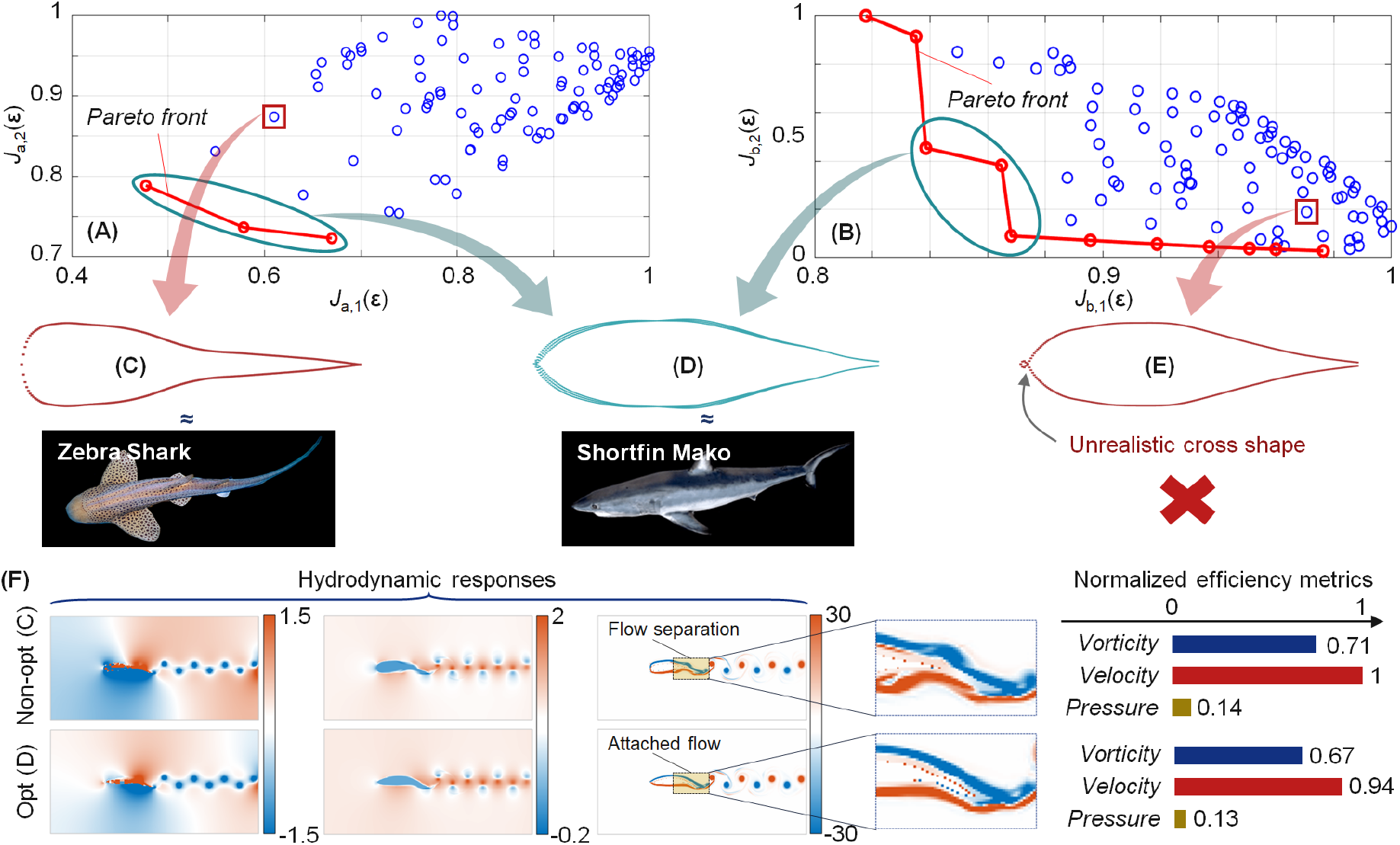
ID-DMD–based multi-objective hydrodynamic design revealing Pareto-optimal trade-offs between competing performance metrics. **(A).** A two-objective Pareto map defined by the normalized dynamic range of vorticity, *J*_*a*,1_(**ε**_d_), and mean vorticity, *J*_*a*,2_(**ε**_d_), where blue circles denote the design space explored by the ID-DMD, while the red curve indicates the Pareto front. **(B)**. A two-objective Pareto map defined by the normalized induced velocity, *J*_*b*,1_(**ε**_d_), and flow field pressure, *J*_*b*,2_(**ε**_d_). **(C)**. A representative non-optimized design lying away from the Pareto fronts, producing a body shape closely resembling a Zebra Shark-type morphology. **(D)**. The fish body shape corresponds to a Pareto-optimal solution shared by both objective spaces (A and B). This geometry exhibits strong similarity to a Shortfin Mako-type morphology. **(E)**. An example of an invalid morphology generated outside the optimization regime, showing a crossed head curve. **(F)** CFD-based validation confirms that the optimized Mako-like morphology facilitates a streamlined, attached flow along the body, significantly outperforming the non-optimized zebra-like profile across all key propulsive efficiency metrics (Lower vorticity range, induced velocity and pressure).

The results demonstrate how engineering solutions can be ex-tended beyond biological observation by systematically exploring the remarkable variation in morphology among fish species, or potentially among other groups of organisms. By unifying morphology, physics, and concepts of locomotor optimization, this framework transforms bio-inspired design from an act of mimicry into a predictive science. Moreover, the methodology reveals a design manifold on which fish geometries are constrained, reinforcing prior work showing how a wide variety of fish phenotypes and levels of locomotor performance have evolved to enable fish to occupy a wide range of ecological niches [51, 52, 53, 22].

## Discussion

### Contributions and Significance

Decomposition of dorsal fish shapes across 15 diverse species reveals a fundamental principle of biological design: the existence of a common fusiform base-line geometry. The remarkable consistency of the first PCA mode across species suggests that, despite the vast morphological diversity observed in nature [54, 55], fishes appear to share a conserved structural foundation for fluid environments. The higher-order modes (Modes 2-4) quantify the evolutionary fine-tuning required for specialized ecological niches. The results demonstrate that the complex morphological space of fishes can be projected into a low-dimensional manifold for both evolutionary biology and bio-inspired engineering research.

By integrating the ID-DMD with morphological analysis, we have moved beyond black-box performance prediction to establish a mechanistic link between shape and hydrodynamics. The modal analysis identifies specific flow-structure interactions that define locomotive efficiencies. These findings provide a physical basis for the long-standing biological hypothesis that tail aspect ratio and curvature are primary drivers of locomotion. This approach has the potential to act as a powerful complement to valuable existing methods to use computational fluid dynamics analysis on fish-like bodies [56, 57], as well as studies of fish kinematics from live specimens [58, 59, 60, 61].

The proposed data-driven framework enables artificial evolutionary exploration at orders of magnitude faster than conventional CFD. In multi-objective optimization, the resulting shapes converged toward the morphology of a high-performance swimmer, the Shortfin Mako shark, demonstrating the biological relevance of the ID-DMD model. This result reinforces prior work demonstrating that natural selection has played a powerful role in shaping the diversity of fish shapes [55], and which continues to play a critical role in bioinspired design for robots [22, 62, 63].

### Scalability Analysis

While these 2D, scale-normalized simulations provide clear hydrodynamic insights, they serve as a methodological proof-of-concept rather than a definitive biological generalization. Real-world aquatic locomotion represents a multidimensional optimization problem involving complex fluid-structure interactions, flexible-body dynamics, and time-varying kinematics within 3D fluid fields.

The extension of this framework to 3D volumetric morphometrics, incorporating intricate caudal and pectoral fin geometries, is demonstrated in Fig.4. Our results indicate that even for high-dimensional, complex geometries, PCA effectively identifies a sparse set of dominant modes governed by a limited number of design parameters (See *Supplementary Information Section V*). The corresponding vortex shedding patterns, shown in Fig.4 (C), illustrate that applying ID-DMD to 3D hydrodynamics is a computationally straightforward operation when performed on these dominant modes. Collectively, these findings validate the proposed morphological design approach for higher-dimensional applications, bridging the gap between simplified 2D contours and high-fidelity, 3D biomimetic engineering.

**Figure 4:**
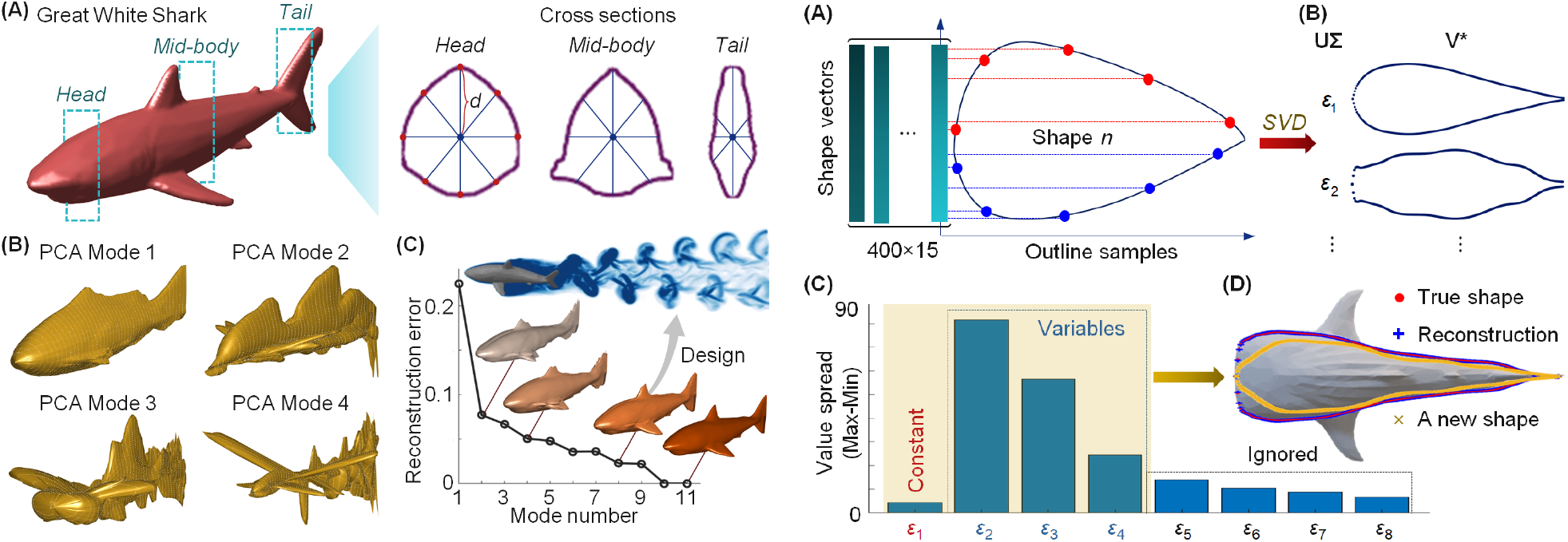
Morphological decomposition and reconstruction of 3D fish body using PCA. **(A).** The 3D model of a Great White Shark in a natural swimming position, showing selected cross-sectional planes at head, mid-body, and caudal peduncle. Detailed cross-sections (right) illustrate the radial distance *d* from the central axis to the body surface. This radial mapping is used for the PCA of the 3D shapes, under the assumption of a convex body profile to simplify the geometric decomposition. **(B)**. Visualization of the first four PCA modes, representing the primary geometric variations captured from the morphological dataset. Each mode highlights different structural components, such as bulk volume, fin placement, and body elongation. **(C)**. Reconstruction error as a function of the number of PCA modes utilized. The plot demonstrates that as the mode number increases, the reconstruction error decreases, allowing for the design of 3D hydrodynamic performances, as shown by the wake turbulence visualization.

### Limitations and Future Works

A critical next step involves delving into the fundamental design principles across broader Reynolds (*Re*) and Strouhal (*St*) regimes. By integrating highfidelity 3D CFD and experimental flume data, we can systematically explore how morphological optimality shifts across different fluid environments. Ultimately, this framework lays the foundation for a digital twin of evolutionary mechanics, providing a powerful tool to decode the subtle hydrodynamic secrets of apex marine predators. Beyond biological discovery, such insights will enable the development of next-generation autonomous underwater vehicles through the application of these bio-inspired design rules.

## Materials and Methods

### Morphological Parameterization

To represent the morphological diversity of fish bodies within a continuous and tractable manifold, we applied a PCA approach to the body shapes of all 15 fish species. For hydrodynamic and morphological analysis, we restricted our study to the two-dimensional body shapes captured from a top-down view. Here, all the fish bodies are scaled to the same length, which means the size effects of the fish are not considered. To enable parameterization, each dorsal-view outline curve was sampled with 400 points clockwise to formulate a shape vector (Fig.5 (A)). All vectors of the 15 fish body shapes are stacked as a shape matrix **B** ∈ ℝ^15*×*400^ (Fig.5 (B)).

**Figure 5:**
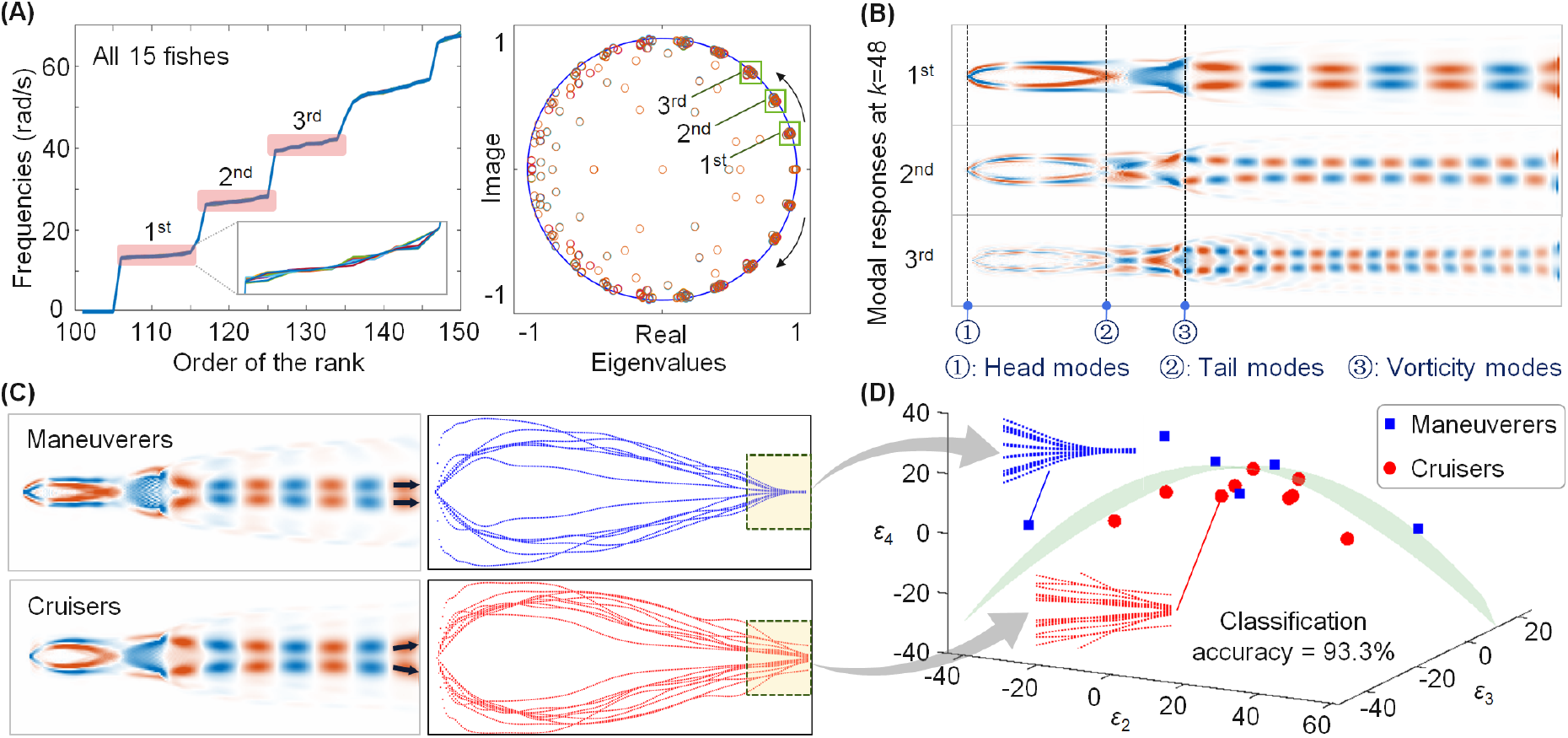
Morphological parameterisation and regeneration of fish body shapes. (**A).** Outline points (400) are sampled along each fish body and combined into a single shape vector per specimen. These shape vectors for 15 fish are stacked into a shape matrix. **(B)**. SVD is applied to identify dominant morphological parameters (*ε*_1_, *ε*_2_, *ε*_3_, *ε*_4_, …) and their associated principal PCA modes, capturing the main geometric variations. **(C)**. The contribution of each morphological parameter is quantified. The first parameter (*ε*_1_) is treated as a constant, *ε*_2_, *ε*_3_, *ε*_4_ are the key design variables controlling body geometry, while higher-order modes are negligible and ignored. **(D)**. Using the reduced set of four shape modes, the approach enables accurate reconstruction of a real fish body (i.e. the Whale Shark) with 3% total reconstruction error, and the generation of new shapes by varying the dominant parameters.

By applying the SVD to the shape matrix **B**, there is

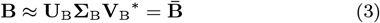

where 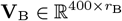 is defined as the PCA modes of the fish bodies, which 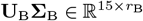 includes the coefficients of the PCA modes (Fig.5 (C)). 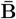 denotes the reconstructed body shapes. *r*_B_≤ 15 is the truncated rank of the shape matrix.

Fig.5 (D) indicate that the first four shape modes with *r*_B_ = 4 capture the dominant morphological features that contribute most significantly to the overall geometry and, consequently, to the hydrodynamic performance. The total reconstruction error of a fish body is defined as

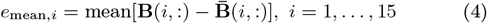

where the shape reconstruction results are shown in *Fig.S1*.

### Comparison with Machine Learning Methods

The ID-DMD, as well as neural operator learning approaches like DeepONet [30] and FNO [31], enable the extraction of swimming hydrodynamics in a fully data-driven manner, without the need for explicit governing equations. In this work, we provide a direct comparison of computational efficiency, accuracy, and interpretability between ID-DMD and these methods in the context of fish hydrodynamics. The comparative results (Bluegill) are summarized in Table.2. The settings of DeepONet and FNO are shown in *Fig.S3*. The results indicate that ID-DMD is still the most efficient and accurate approach for fish body hydrodynamic design, while also providing clear physical interpretability. Although the predicted flow fields and reconstructed initial conditions still exhibit residual errors, the method provides a valuable tool for preliminary design exploration and for guiding subsequent high-fidelity optimization.

**Table 2:** Comparison of advanced operator learning approaches^*1^.

|  | ID-DMD | DeepONet | FNO |
| --- | --- | --- | --- |
| Training time | 14.8 s (CPU) | 0.37 h (GPU) | 16.22 h (GPU) |
| Simulation time | 0.02 s (CPU) | 0.15 s (GPU) | 1.2 s (GPU) |
| Mean relative error (Max)*2 | $3.2 \times 10^{-3}$ | $2.61 \times 10^{-2}$ | $2.45 \times 10^{-2}$ |
| Interpretability | Yes | No | No |

\*1 The Google Colab A100 GPU is applied for comparison.
\*2 The maximum mean relative error is $\max_{k=1, \dots, 48} [\text{mean}(\eta_k)]$

### Mode-based Morphological Analysis

The ID-DMD framework not only provides accurate predictions of the hydrodynamic fields around diverse fish body shapes, but also extracts dynamic modes with clear physical interpretability, offering new insights for morphological analysis. The modal frequencies derived from the ID-DMD representation are symmetric about zero, with the positive frequencies for all 15 fish species displayed in Fig.6 (A). Notably, the first-order mode consistently captures the dominant tail-beat frequency (*ω*_1_ ≈ 13.2 rad/s), which characterizes the primary oscillatory motion of the caudal fin during steady swimming (*ω*_s_ = 4*π* rad/s). This mode reflects the fundamental frequency of propulsion and serves as a key dynamic signature of each species’ locomotor pattern.

**Figure 6:**
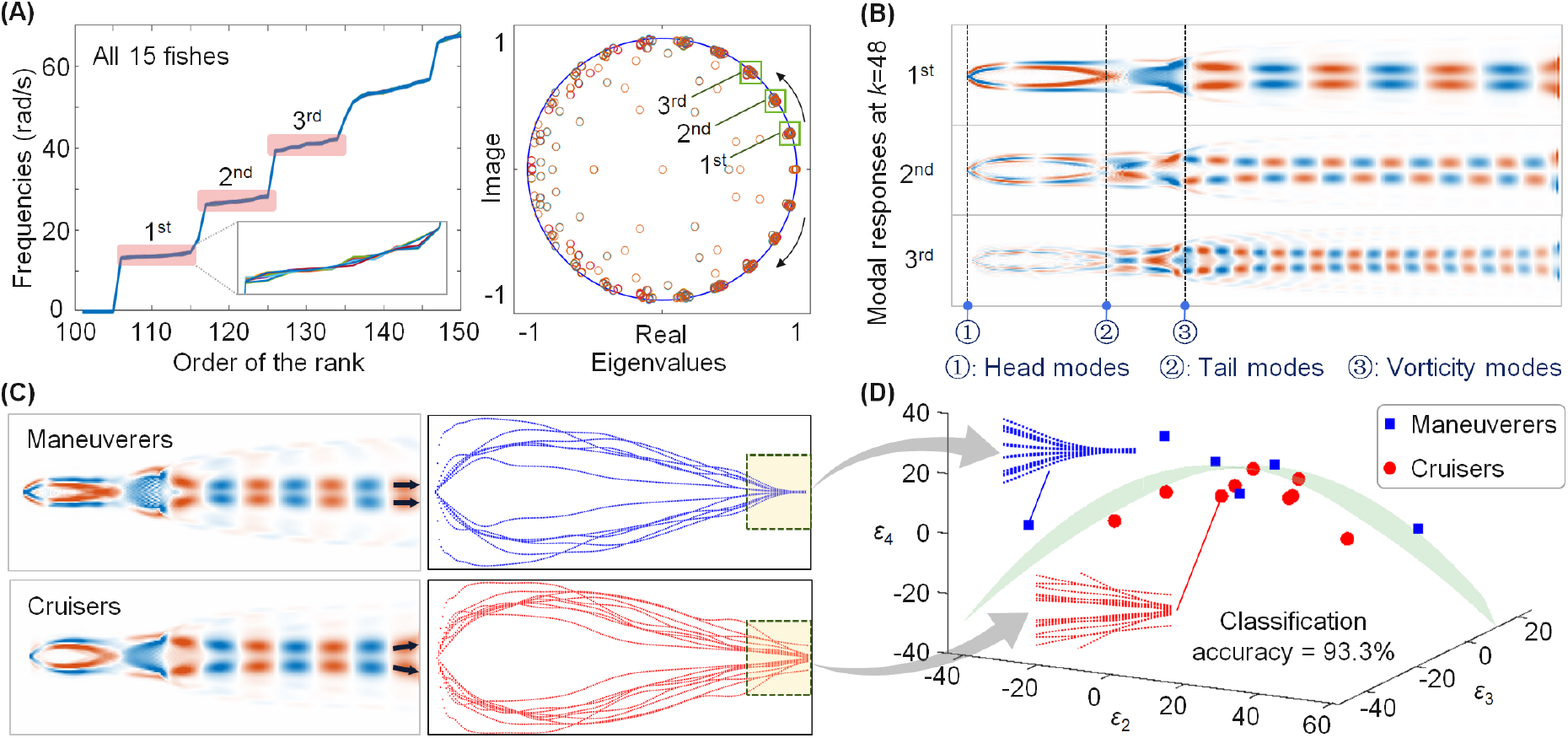
Modal analysis of fish body-fluid dynamic interaction. **(A).** Modal frequencies of local fish body motion obtained from the ID-DMD, showing a clear separation between the first-, second-, and third-order dominant modes. **(B)**. Representative modal responses at *k* = 48, illustrating how different dynamical modes correspond to distinct patterns of body deformation and wake fields. **(C)**. Slender or broader tail shapes result in different types of modal responses. Slender tail configuration tends to concentrate trailing-edge vortices and is often associated with flexibility or high thrust efficiency during burst and rapid swimming, known as *Maneuverers*, including Great White Sharks, Epaulette Shark, Largemouth Bass, Sand Tiger Shark, Striped Bass, Whale Shark; Broader tail configuration facilitates broader vortex shedding, supporting sustained, energy-efficient propulsion during cruising, known as *Cruisers*, including Atlantic Sturgeon, Blue Gill, Bronze Whaler Shark, Leopard Catshark, Puffadder Shyshark, Sevengill Shark, Shortfin Mako, Zebra Shark. **(D)**. Classification of fish body shapes using PCA Modes for the characterization of fish hydrodynamics. The two fish groups are clearly distinguished by the principal PCA modes, demonstrating a functional pipeline where mode coefficients dictate body morphology, which in turn determines the hydrodynamic performance required for specific locomotory functions. The Largemouth Bass is misclassified due to the truncation errors in its morphological reconstruction.

These modal responses (*Supplementary Information Section III*) clearly reveal the characteristic oscillation patterns of the fish head and tail, as well as the associated vorticity fields, as illustrated in Fig.6 (B). These results reveal a consistent dynamic behavior across species:

- The head remains relatively stable;
- The tail undergoes periodic, pendulum-like swinging;
- The traveling vortices exhibit a regular and periodic shedding pattern that reflects the underlying propulsion mechanism.

Together, they highlight the coordinated interplay between body kinematics and fluid dynamics during steady swimming.

In Fig.6 (C), the “upward-bending” response observed in slender-tailed species (e.g., Great White Sharks) suggests flexibility or high propulsive efficiency during burst and rapid swimming. In contrast, “horizontally aligned” responses are found in broader-tailed species (e.g., Shortfin Mako), supporting sustained, energy-efficient propulsion during cruising (See details in *Supplementary Information Section III*).

### Inverse Design for Hydrodynamics

The hydrodynamic design of dorsal fish body shapes can be formulated as

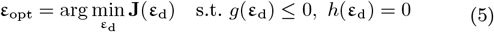

where **J**(**ε**_d_) denotes a set of objective (loss) functions *J*_1_(**ε**_d_), *J*_2_(**ε**_d_), … for the optimal design problem, while *g*(**ε**_d_) and *h*(**ε**_d_) represent the inequality and equality constraints, respectively.

Standard optimization algorithms, such as Gradient Descent methods [64] and Genetic Algorithms [65], can then be employed to solve the design problem, enabling the hydrodynamic optimization of fish body shapes.

In this study, the following objective functions are considered for the design of fish body shape under the constraint of

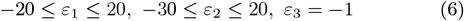

1. The dynamic range of vorticity (Δ*V*):

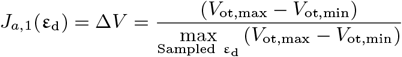

where *V*_ot_ is the vorticity and *J*_*a*,1_(**ε**_d_) measures the normalized range of vorticity, where a minimized range indicates a reduction in flow separation and excessive shear layer intensity.
2. The mean value of vorticity:

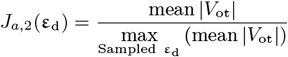

Minimizing the vorticity range and mean magnitude ensures a smooth vorticity signature. This characteristic is a hallmark of high propulsive economy, as it indicates that the body suppresses high-energy vortex shedding that would otherwise dissipate as heat [66, 67].
3. The normalized induced velocity:

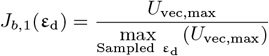

where *U*_vec_ is the induced velocity of the fish. Minimizing the induced velocity relative to the swimming speed is critical to reducing the kinetic energy wasted in the wake [68, 69, 70].
4. The normalized pressure of the flow field:

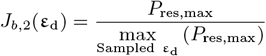

where *P*_res_ is the pressure of the flow field. Minimizing the pressure dynamic range is key to maintaining a favorable pressure gradient. This minimizes pressure drag by preventing boundary layer separation [71, 72].

By achieving a simultaneous reduction in vorticity, induced velocity and pressure fluctuations, the optimization identifies a morphology that maximizes propulsive efficiency. This ensures a streamlined, low-dissipation interaction with the fluid, consistent with the specialized physiology of apex predators like the Shortfin Mako.

### Evaluation of Initial Flows

In designing the hydrodynamics of fish bodies, it is essential to evaluate the initial conditions corresponding to various design parameters. The initial flow fields were reconstructed via PCA-based characteristic parameters predicted directly from morphological design variables (*Fig.S4*). The high predictive accuracy of this linear mapping ensures a robust starting point for subsequent DMD analysis. See *Supplementary Information Section IV* for further details.

## Acknowledgments

YPZ acknowledges support from the Queen Mary University of London Startup Funding (SEM9307B). We also thank Hailong Liu and Sijie Wang at the University of Groningen for their contributions to the CFD simulations and operator-learning comparisons. JNK acknowledges support from the Air Force Office of Scientific Research (FA9550-24-1-0141).

## Supporting Information

### Section I: Morphological analysis of dorsal fish bodies

The dataset employed in this study is derived from the Digital Life Project (www.digitallife3d.org/3d-model), developed at the University of Massachusetts Amherst, which offers high-resolution 3D scans of 15 living fish species listed in **Table.1**. Each 3D model is processed to extract the dorsal outline, from which 400 points are uniformly sampled along the body contour to form a high-resolution 2D shape profile. All sampling points are stacked as a shape matrix **B** ∈ ℝ^15*×*400^ (**Fig.4**).

In the parameterization of fish bodies, the PCA allows complex fish body geometries to be represented using only a few dominant shape modes. These modes are ranked by their energy content, which is determined by the magnitude of their corresponding eigen-values. By applying the SVD to the shape matrix **B**, there is [73]

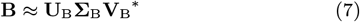

where 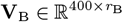 is defined as the PCA modes of the fish bodies, which 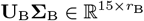 includes the coefficients of the PCA modes. *r*_B_ ≤15 is the truncated rank of the shape matrix.

Specifically, the PCA coefficients are represented as

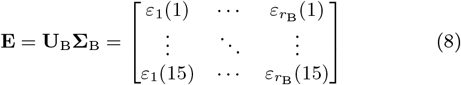

An important feature of the PCA-based representation is its ability to generate new fish body shapes by systematically varying the PCA coefficients 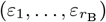. Here, higher-order PCA modes capture increasingly fine-scale geometric details, while lower-order modes encode the most dominant features of the shape.

The first 4 PCA modes and their coefficients across all 15 species are shown in **Fig.7 (A)**, where the coefficient deviations of the *r*th order PCA are calculated by removing the mean values of *ε*_*r*_ over all 15 species. It can be observed that the first PCA mode and its associated coefficients are constant *ε*_1_(*i*) ≈ − 1.96 *×*10^3^ for *i* = 1, …, 15, which reveals a common underlying fusiform body shape across the fish [74].

**Figure 7:**
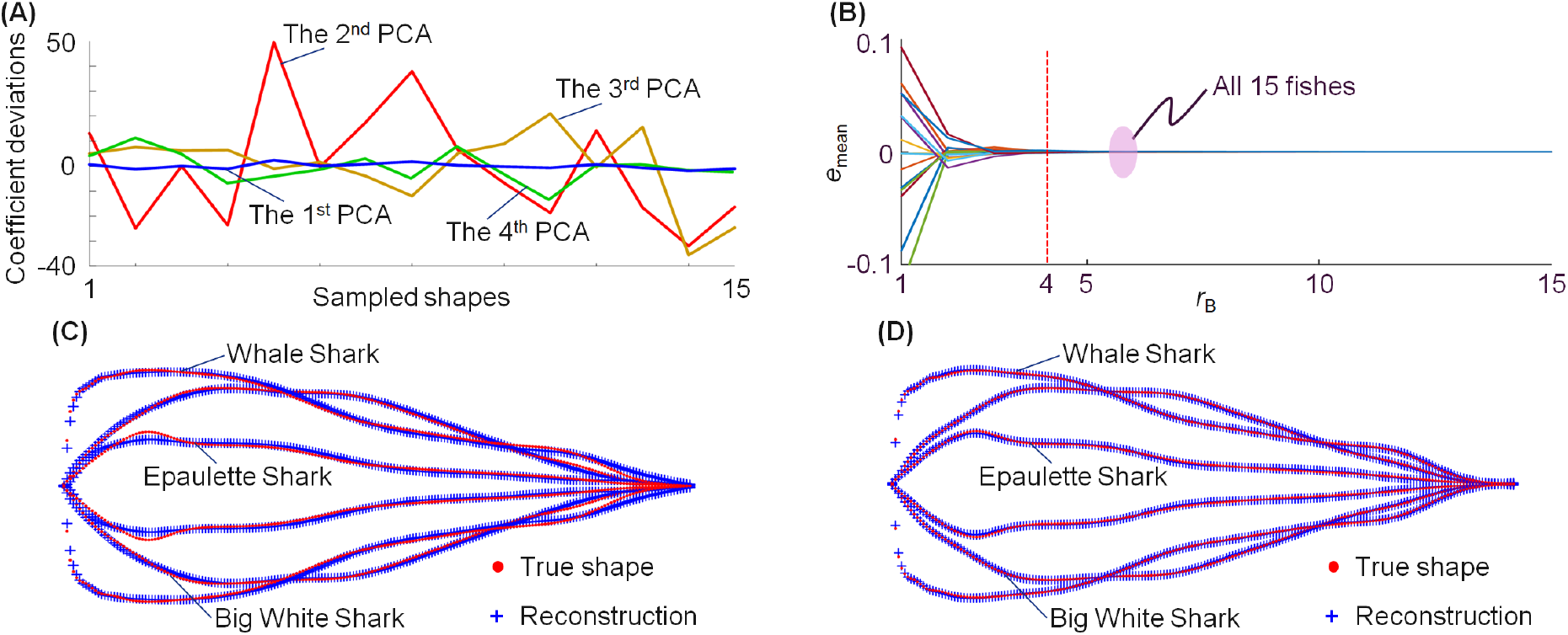
Reconstruction of fish body shapes. **(A).** Variation in PCA coefficients for all 15 fish bodies. The first coefficient remains nearly constant across species, indicating a shared baseline morphology primarily captured by the first mode. **(B)**. Mean reconstruction error as a function of the number of retained PCA modes. The error rapidly decreases and approaches zero with four modes, showing that the dominant morphological features are captured by a low-dimensional representation. **(C)**. Reconstruction of fish body shapes using the first four PCA modes. **(D)** Reconstruction of fish body shapes using the first eight PCA modes.

The error curve in **Fig.7 (B)** shows a rapid convergence within the first 4 modes, where the reconstruction errors for each fish body shape are evaluated in equation [4].

Therefore, the coefficients of the second, third, and fourth PCA modes are then treated as 3 free design parameters. **Fig.7 (C)** and **(D)** demonstrate the reconstruction of fish body shapes based on the first *r*_B_ = 4 and *r*_B_ = 8 shape modes. It is clear that the first 4 modes capture the dominant morphological features that contribute most significantly to the overall geometry, while higher-order modes are omitted due to their limited impact on the dorsal fish body shape reconstruction.

### Section II: ID-DMD for fish hydrodynamics

#### CFD Simulation of Fish Bodies

Hydrodynamics around 2D fish bodies are evaluated via the CFD simulation. The CFD simulation platform is the WaterLily, a differentiable and backend-agnostic Julia solver [75]. As illustrated in **Fig.8**, the 2D flow field is set with a length of 4.0 fish Body Length (BL), a width of 2.0 BL, and the Reynolds number is *R*_e_ = 1e^5^

**Figure 8:**
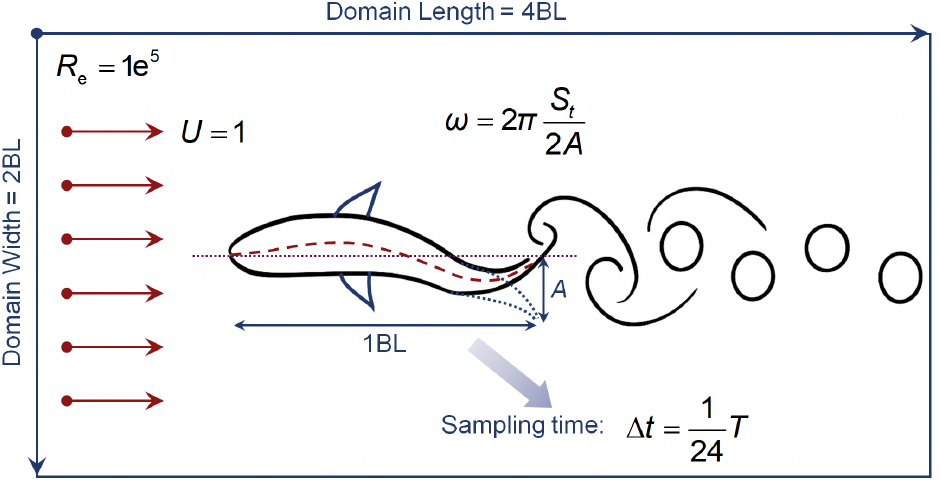
Set up of the CFD simulation of fish bodies.

Here, the WaterLily employs the Cartesian-mesh method using constant spacing. The characteristic water flow applies to the unit horizontal inlet velocity. The upper and lower walls are impermeable with free-slip tangential conditions. The swinging undulation amplitude of the 2D fish is set to 0.1 BL. Once the undulating flow field reaches a steady state, a total of 2 undulation periods are measured with the motion frequency in physical time is given by

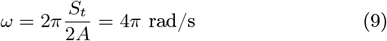

where *S*_*t*_ = 0.4 is the Strouhal number representing a representative intensive propulsive state, and *A* = 0.1 is the Amplitude-to-BL ratio.

The sampling time is Δ*t* = *T/*24 of an undulation period of *T* = 2*A/S*_*t*_ = 0.5 s. In this study, the Vortex dynamics, pressure distribution, and induced velocity field data are extracted from the simulated flow domain.

#### ID-DMD for the hydrodynamics of fish

The ID-DMD approach offers a lightweight, fully data-driven framework for modelling and prediction in digital system design [36].

Let **x** _*k*_∈ ℝ^*m*^ with *m* being the number of states denote the state vectors of a system (e.g., vorticity, velocity, and pressure) at a discrete time step *k*. For fluid dynamics [40], these vectors can be obtained by flattening the system snapshots at times *k*Δ*t* on a pixel level, where Δ*t* is the sampling time. Let **ε**_d_ = {*ε*_2_, *ε*_3_, *ε*_4_}represent the design parameters of fish bodies. Then, the ID-DMD representation is defined as

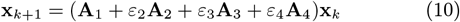

where **A**_*i*_, *i* = 1, 2, 3, 4 are operator matrices with constant entries.

To identify the ID-DMD model of fish hydrodynamics, we conducted CFD simulations on 9 representative fish body shapes. Their resulting flow vorticities are used as training data. The remaining 6 species (Great White Shark, Bluegill, Leopard Catshark, Largemouth Bass, Shortfin Mako, and Striped Bass) are reserved for testing and evaluation to assess the model’s predictive capability.

For each fish shape, the simulation generated 48 time-resolved snapshots over the 4 tail-swinging cycles. Each snapshot consists of spatial fields with a resolution of 386*×* 194 pixels, covering the region around the fish body. Each snapshot of the vorticity was reshaped into a column vectors **x**_1_, …, **x**_48_ and stacked to form the high-dimensional state matrices

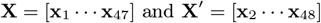

The ID-DMD can be represented as

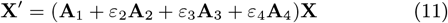

The objective of the ID-DMD approach is to directly evaluate the eigenfrequencies *s*_*j*_ and eigenvectors **w**_*j*_ for *j* ∈ ℤ^+^ of the parameterized linear operator

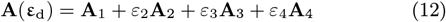

Therefore, for any given set of design parameters **ε** = {*ε*_2_, *ε*_3_, *ε*_4_}, the system state at time step *k* can then be obtained as:

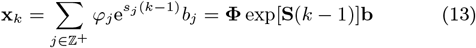

where **b** = **Φ**^†^**x**_1_ and “†” denotes the pseudo-inverse, **x**_1_ represents the initial state of the system. Each eigenfrequency *s*_*j*_ = *σ*_*j*_ + j*ω*_*j*_ involves the frequency *ω*_*j*_ and the decay rate *σ*_*j*_.

#### The ID-DMD Algorithm

The advantage of the ID-DMD algorithm (Algorithm 1) is to evaluate these eigenvalues and eigenvectors is via a reduced order operator.

To account for multiple parameter sets in training, we collect all data into global regression and regressor matrices. Let

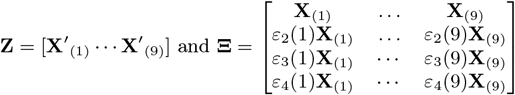

where **X**_(*l*)_ and **X**′_(*l*)_ are data matrices from the *l*th parameter set. This yields the compact form

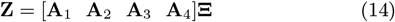

The ID-DMD algorithm is illustrated in Algorithm.1, where Here,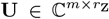 and 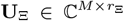 are unitary matrices truncated to ranks *r*_Z_ ≤ *m* and *r*_Ξ_ ≤*m*, respectively, with *M* being the number of rows in **Ξ**.

*Remark 1*. In physics, a complex frequency with a positive real part often indicates an unstable system that diverges over time. To ensure stability, we enforce that the real part of the frequency satisfies *σ*_*j*_ = 0 if *σ*_*j*_ *>* 0.

*Remark 2*. In the ID-DMD, *r*_*Z*_ and *r*_Ξ_ are two hyperparameters to be defined, representing the truncation orders of **U** and **U**_Ξ_, respectively. These are usually determined based on the cumulative energy of the eigenvalues or iteratively tuned according to the testing performance. Here, we set *r*_*Z*_ = *r*_Ξ_ for the convenience of illustrating the modelling process.

#### Validation of Results

In this study, the truncation ranks were set to *r*_*Z*_ = *r*_Ξ_ = 200. These rank values were selected based on the cumulative energy content of the singular values, ensuring that over 90% of the system dynamics are retained in the reduced-order model.

For any given set of design parameters **ε**_d_ = {*ε*_2_, *ε*_3_, *ε*_4_}, the eigenfrequencies *s*_*j*_ and corresponding eigenvectors ***φ***_*j*_ for *j* = 1, …, 200 can be efficiently computed. Therefore, the hydrodynamic response of a 2D dorsal fish body can be accurately predicted using the modal composition rule in equation [14].

##### Algorithm 1 The ID-DMD algorithm

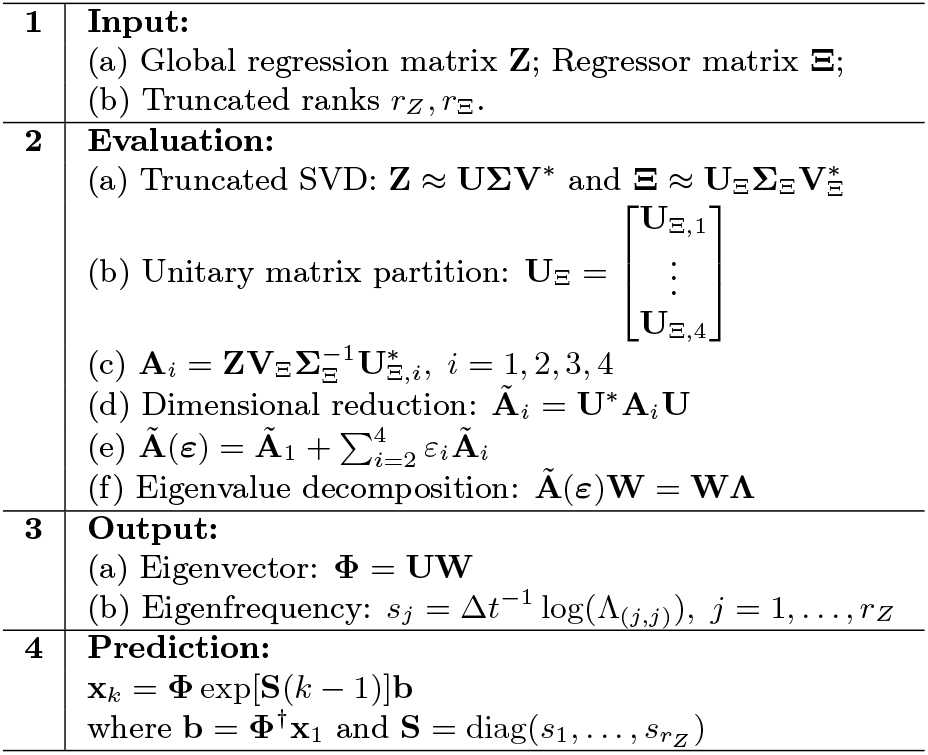

The predictive error is measured by the average of the relative errors over the flow field as “mean(**η**_*k*_)”. The mean and standard deviation of predictive errors of all 6 fish bodies for testing over time are shown in **Table.3**.

**Table 3:**
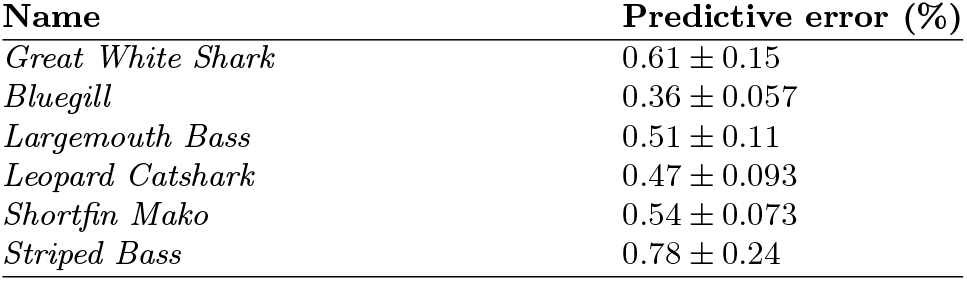
Predictive errors of the 6 fishes for testing.

| Name | Predictive error (%) |
| --- | --- |
| *Great White Shark* | $0.61 \pm 0.15$ |
| *Bluegill* | $0.36 \pm 0.057$ |
| *Largemouth Bass* | $0.51 \pm 0.11$ |
| *Leopard Catshark* | $0.47 \pm 0.093$ |
| *Shortfin Mako* | $0.54 \pm 0.073$ |
| *Striped Bass* | $0.78 \pm 0.24$ |

#### Comparison Results

The extended architectures of both the parametric Vanilla MIONet (DeepONet) and Fourier-MIONet (FNO), adapted to this dataset structure, are illustrated in **Fig.9** To ensure a fair comparison with the proposed ID-DMD framework, two multi-input operator learning models were trained on the same six-dimensional dataset

**Figure 9:**
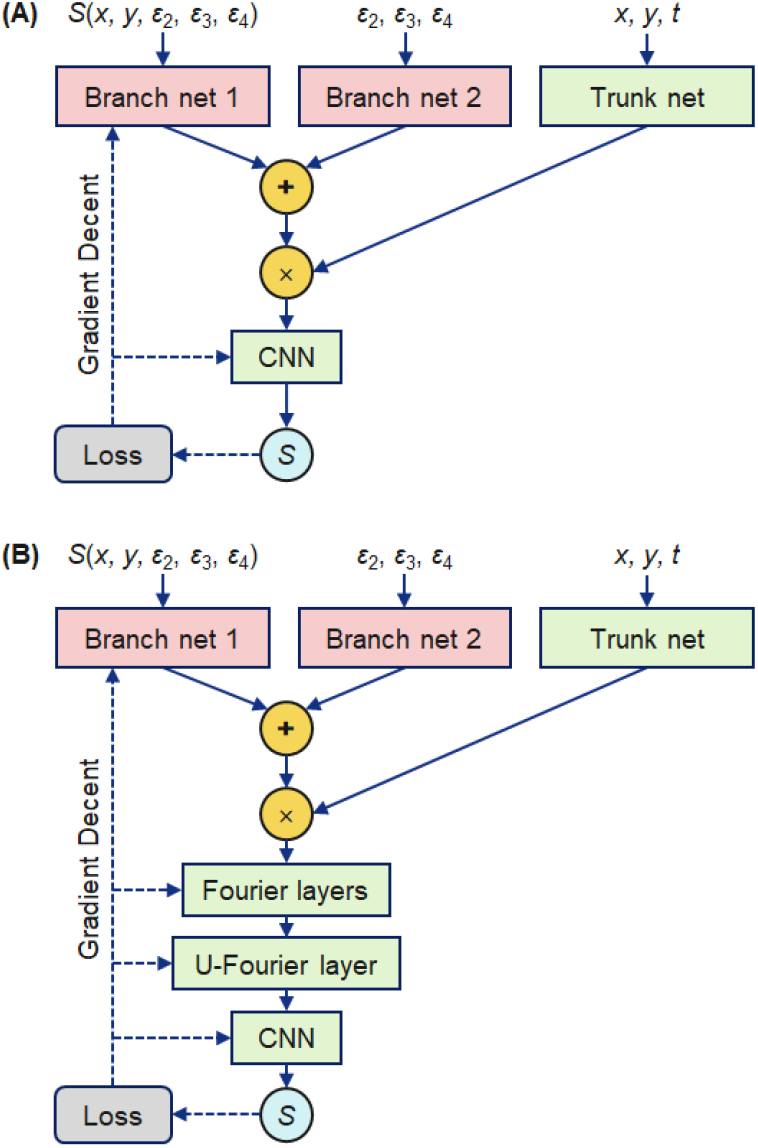
Extended architectures of parametric operator learning networks. **(A).** The architecture of the Vanilla MIONet (Deep-ONet). **(B)**. The architecture of the the Fourier-MIONet (FNO).

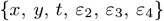

where the spatial grid spans *x* ∈ [0, 3.95], *y* ∈ [0, 1.93], and the time interval is *t* ∈ [0, 0.47].

The parameters *ε*_2_, *ε*_3_, and *ε*_4_ denote the design variables. The Initial Condition (IC) at *t* = 0 is represented by a tensor *S*(*x, y, ε*_2_, *ε*_3_, *ε*_4_), where *S* denotes the flow field.

The vanilla MIONet adopts a triple-network architecture. Two branch networks encode the initial condition *S*(*x, y, ε*_2_, *ε*_3_, *ε*_4_) and the parameters (*ε*_2_, *ε*_3_, *ε*_4_), while a trunk network processes the spatio-temporal inputs (*x, y, t*). Their outputs are combined through element-wise multiplication followed by a linear projection. The model (7,006,145 parameters) is trained for 112,500 epochs using Adam (initial learning rate 0.001, decaying by a factor of 0.9 every 3,375 epochs) with the same loss function described in [76].

The Fourier-MIONet maintains the same input structure but introduces three modifications: 1) shallower networks for improved efficiency, 2) element-wise addition for feature fusion, and 3) a decoder consisting of three Fourier layers, one U-Fourier layer, and one linear layer to better capture spectral dynamics. With 1,695,453 trainable parameters, it is trained for 101,250 epochs with a gradually decaying learning rate starting from 0.001 as described in [77].

The Fourier decoder enhances the model’s ability to represent complex spatio-temporal patterns compared with the vanilla architecture.

### Section III: Interpretability of the ID-DMD model

The ID-DMD framework not only provides accurate predictions of the hydrodynamic fields around diverse fish body shapes, but also extracts dynamic modes with clear physical interpretability, offering new insights for morphological analysis. While many studies have examined the relationship between fish morphology and hydrodynamic performance [78, 79, 80], we present a unified, data-driven approach that directly links geometric features to flow dynamics. Leveraging this capability, we reveal how specific tail shapes govern vortex formation, facilitating both scientific reasoning about evolutionary adaptations and the design of bio-inspired systems.

To investigate the morphological logic underlying dominant dynamic modes, the dorsal fish body and flow responses reconstructed using the first, second, and third order modes are computed as below.

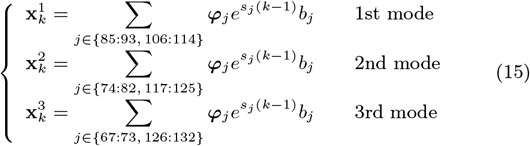

These modal responses clearly reveal the characteristic oscillation patterns of the fish head and tail, as well as the associated vorticity fields, as illustrated in **Fig.5**.

By exploring the first-order modal responses, we can observe two types of response modes as shown in **Fig.5**. The two response modes are categorized as the “Upward-bending mode” and the “Horizontally aligned mode.” Interestingly, all species with the Upward-bending mode (∼ 10^*°*^) display a long and thin tail configuration. This morphology tends to concentrate trailing-edge vortices and is often associated with flexibility or high thrust efficiency during burst and rapid swimming (The maneuverers [81]). In contrast, the Horizontally aligned mode is consistently linked to a short and broad tail configuration. This structure facilitates broader vortex shedding, typically supporting sustained, energy-efficient propulsion during cruising (the cruisers [82]). These mode distinctions reveal a clear coupling between body dynamics, tail morphology, and swimming strategy.

The ID-DMD-based modal analysis provides a powerful tool to link fish body shape with swimming dynamics. It captures both dominant and higher-order flow patterns in a physically interpretable way, helping to explain how different morphologies affect propulsion. This offers new insights into fish evolution and supports the design of efficient bio-inspired swimming systems.

It is worth pointing out that while the proposed method reveals clear relationships between hydrodynamic responses and fish body shapes, it does not fully explain the underlying biological behaviors and evolutionary strategies of fish. Addressing these challenges will require further interdisciplinary research that integrates insights from biology, biomechanics, and fluid dynamics [83, 84, 85].

### Section IV: The ID-DMD model based inverse design

#### Evaluation of Initial Conditions

In designing the hydrodynamics of fish bodies, it is essential to evaluate the initial conditions corresponding to various design parameters, denoted as **x**_1_ ∈ℝ^*m*^. Given the high dimensionality of **x**_1_, a reduced-order approach is required for efficient evaluation. The proposed method is illustrated in **Fig.10** and described below.

**Figure 10:**
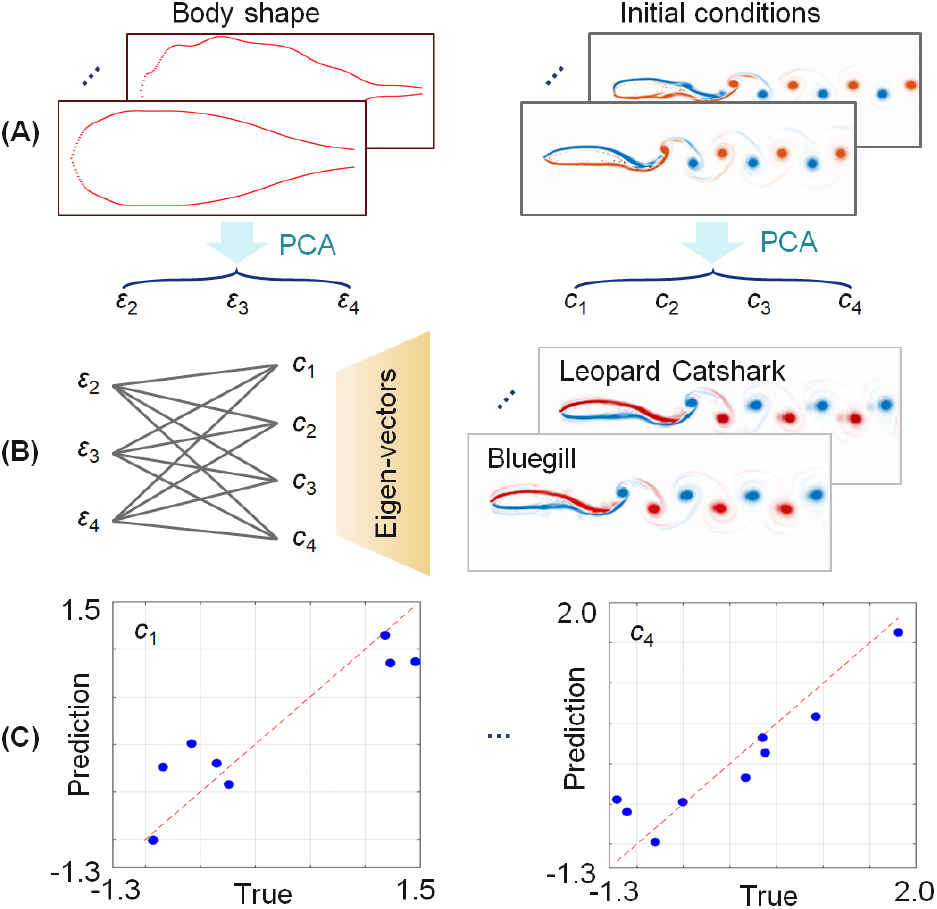
Prediction of initial flow conditions for fish-body hydrodynamics in latent space. **(A).** PCA decomposition of dorsal body shapes and the corresponding initial flow fields. **(B)**. Mapping from shape parameters to the PCA coefficients of the initial conditions, enabling reconstruction of flow fields for unseen fish designs. Examples for Bluegill and Leopard Catshark are shown. **(C)**. Comparison between predicted and true PCA coefficients for representative modes.

First, PCA of both the 2D dorsal fish body outlines and their corresponding initial flow fields are conducted using singular value decomposition (SVD), as illustrated in **Fig.4**, where the design parameters **ε**_d_ = {*ε*_2_, *ε*_3_, *ε*_4_}correspond to the second, third, and fourth PCA coefficients of the body shape.

The initial flow fields are reshaped into high-dimensional vectors, and the initial-condition matrix constructed from the training data (identical to the training set used for ID-DMD) is defined as

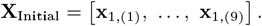

The singular value decomposition (SVD) of the initial-condition matrix is obtained as

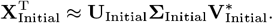

Here the truncated rank is *r*_Initial_ = 4. The matrix **V**_Initial_∈ ℝ^*m×*4^ contains the PCA modes of the initial conditions, while **U**_Initial_ **∑**_Initial_ ℝ^9*×*4^ contains the coefficients of the PCA modes, denoted as **c** = {*c*_1_, *c*_2_, *c*_3_, *c*_4_}.

The design parameters *ε*_2_, *ε*_3_, and *ε*_4_ are then mapped to the PCA coefficients *c*_1_, *c*_2_, *c*_3_, and *c*_4_ via polynomial functions, as illustrated in **Fig.10 (B)**:

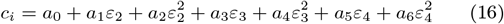

for *i* = 1, 2, 3, 4, where *a*_0_, …, *a*_6_ are the polynomial coefficients.

The prediction results are shown in **Fig.10 (C)**. The initial conditions corresponding to different design parameters are finally reconstructed using the PCA modes **V**_Initial_.

